# Mapping Fusarium Wilt and Sterility Mosaic Disease Resistance-Associated Genomic Regions and Haplotype Variants in Pigeonpea

**DOI:** 10.1101/2025.06.19.660531

**Authors:** Sagar Krushnaji Rangari, Suma Krishnappa, Mamta Sharma, Vinay K Sharma, Namita Dube, Sunil S Gangurde, Vinay Sharma, Ragavendran Abbai, Naresh Bomma, Shruthi Belliappa, Karma L Bhutia, Mahendar Thudi, Prakash I Gangashetty, Manish K Pandey

## Abstract

The occurrence of fusarium wilt and sterility mosaic disease in genial conditions causes significant yield losses in pigeonpea. The present genome-wide association study (GWAS) was employed on 176-genotype panel to identify candidate genomic regions associated with FW and SMD resistance in pigeonpea. A total of 869,447 filtered SNP markers were used for GWAS analysis. GWAS analysis identified significant genomic regions for FW on chromosomes 05, 10, and 11 and for SMD on chromosome 09. Four resistant sources, namely, ICP20096, ICP20097, ICPL87119 and ICP13304 were identified as a resistance source for both FW and SMD. Total 12 markers from chromosome 11 for FW and 4 markers from chromosome 09 for SMD were validated using the KASP genotyping approach. We have identified resistant haplotypes for FW on chromosome 10 and 11 and for SMD on chromosome 09. Moreover, we have identified four InDels for FW on chromosome 5 in *Two component response regulator gene* (*Cc_11098*) and four InDels for SMD on chromosome 9 in genes, namely, *LRR and NB-ARC domain disease resistance protein* (*Cc_21069*), *Flavonoid 3’, 5’-hydroxylase* (*Cc_21070*) respectively. For FW, *Cc_10507586* region from chromosome 5 is identified in ICPL87119 and ICPL20096 donors. A region *Cc_21637656* from chromosome 11 is identified in the ICPL87119 but not in the ICPL20096. Similarly, a region *Cc_4438493* from chromosome 10 is identified in the ICPL20096 but not in the ICPL87119. For SMD, *Cc_24131177* region is identified in both the SMD resistance donors (ICPL20096 and ICPL87119). *Ccsmd04* region from chromosome 4 has been identified in ICPL87119 but not in the ICPL20096. These genomic regions can be brought together into single elite genetic background with the help of the markers identified in this study. The validated markers and donor lines in this study offer valuable tools and resources for the genetic improvement of pigeonpea cultivars with enhanced FW and SMD resistance.

## 1. Introduction

Pigeonpea (*Cajanus cajan* (L.) Millspaugh) is an often-cross pollinated perennial shrub that is traditionally cultivated in semi-arid regions of the world. ∼3,500 years ago, the earlier domestication was initiated in central India from its wild progenitor, *Cajanus cajanifolius* [1]. India is considered as the primary centre of origin for pigeonpea. It is considered the sixth most important legume, which contributes to 5% of the total pulse production (5.32 MT) from the 6.3 Mha with a productivity of 883 kg/ha in the world. It is cultivated mostly in Asia, Africa, Latin America, and the Caribbean region. Among pigeonpea-growing counties, India contributes to ∼79% global annual production [2–3].

Among different production constraints, Fusarium wilt (FW; caused through soil-borne fungus *Fusarium udum* Butler) and Sterility mosaic disease (SMD; caused through pigeonpea sterility mosaic virus) are major in pigeonpea. FW is the most serious biotic threat to pigeonpea production throughout the world. *F. udum* Butler enters the pigeonpea plant through roots and starts infecting seedlings; significant damage is caused at flowering and podding stages. The occurrence of infection has been recorded up to 60%, particularly at the flowering and crop maturity stages of plants [4]; nevertheless, 100% yield losses were also reported in susceptible cultivars [5]. Interveinal chlorosis, internal browning of xylem vessels, loss of turgidity, yellowing and drooping of leaves, and withering and wilting of plants are key symptoms of FW. Total nine variants of *F. udhum* distributed throughout India. Among these, 2 and 3 variants are found most predominant in most pigeonpea-growing areas [6].

For the first time, SMD was identified in 1931 at Pusa, Bihar, and becomes the endemic for the cultivated regions of India and its nearby countries. Approximately over US$ 300 million in yield losses were reported in India alone due to SMD [7]. The characterization of the SMD can be done by observing bushy and stunted plants, whose leaves have been reduced in size with chlorosis on them and lowered in the production of flowers [8]. There are two variants of SMD virus that were observed: one is pigeonpea sterility mosaic virus 1 (PPSMV-1) having five negative-sense RNA segments, and another is pigeonpea sterility mosaic virus −2 (PPSMV-2) having six negative-sense RNA segments [9]. Some cultivated and wild species of pigeonpea were found to support a vector of this virus, *Aceria cajani,* an eriophyid mite. This scenario was only found in the infected plant, which shows the strong relation between these infected plants and virus-carrying vectors [10]. Many popular cultivars such as ICP8863 (Maruti) and JKM-189 are susceptible to SMD and becoming a host to spread this disease in the nearby region.

Although efforts are being made to develop FW and SMD resistant cultivars using conventional breeding, understanding the genetic mechanism and identification of candidate genes will be a potential solution to come up with strategies to breed for resistance. In past two RAPD markers linked two recessive alleles of FW resistance genes were reported [11]. Using Sanger-based bacterial artificial chromosome end sequences and a genetic map, 40.46% pseudomolecules of the pigeonpea genome comprising 29,482 genes have been reported in version 1 [12]. The updated version is also now available with 91.35% pseudomolecules and 48,680 genes [13]. Earlier, in pigeonpea, QTL-seq was used to identify genomic regions for FW and SMD resistance in RIL population ICPL20096 × ICPL332 but no success [14]. Due to use of common resistant and susceptible bulks for both FW and SMD hindering with QTL detection. The same dataset was then used to identify 26 genome-wide InDels affecting 26 candidate genes [15]. Again, three populations [ICPL20096 × ICPL332, ICPL20097 × ICP8863, and ICP8863 × ICPL87119 (F_2_) population] were used to map genomic regions for SMD resistance using genotyping-by-sequencing (GBS) [16]. Among these, two major QTL regions were identified in ICP8863 × ICPL87119 on chromosome Cc03 (*qSMD3.1*) with 34.3% PVE and LOD 2.8, and on chromosome Cc11 (*qSMD11.3*) with 24.2% PVE and LOD 5.8. All these studies were performed using an older version of ‘Asha’ pigeonpea reference genome v1.0, assembled using short reads sequencing [12]. Interestingly, in our GWAS study, we have used the improved telomere-to-telomere ‘Asha’ reference genome v2.0 assembled using Hi-Fi sequencing in combination with Hi-C (High throughput chromosome capture) [13].

On the other hand, genome-wide association studies (GWAS) have become the highly important tool for mapping the regions associated with the variety of plant traits. GWAS has identified marker-trait associations for various traits almost in all the major crops. Multiple models for GWAS are now available, which can even handle multi-locus mapping with improved computational speed and higher power [17]. Post-GWAS analysis of the GWAS results is also becoming important to validate the identified MTAs and candidate genes. The GAPIT package of RStudio provides phenotypic variation explained (PVE) percentage, prediction values such as BLUP (to normalise the data), allele distribution plots for significant markers, and other important figures other than Manhattan and QQ plots. All these outcomes represent GWAS as a powerful tool for mapping the traits. The identification of associated regions, candidate genes, and diagnostic markers are highly necessary to improve the traits and develop climate-resilient and high-yielding crops to fulfil the requirements of the world’s growing population.

In view of the above, the present study identified the novel SNP and Indel markers, candidate genes, and haplotypes associated with the FW and SMD resistance using the updated genome version of pigeonpea. It has also been shown that the genes identified in this study are involved in the disease-resistance-related pathways. Moreover, the KASP based markers validation was done for identified regions.

## 2. Material and methods

### 2.1 Plant material and field design

For GWAS analysis we have selected 176 genotypes from the reference set based on the availability of sequencing data. Among 176 genotypes, two are from Australia, one from Bangladesh, one from China, one from Ghana, one from Cambodia, 30 from ICRISAT, 106 from various parts of India, two from Italy, one from Jamaica, one from Kenya, two from Myanmar, five from Nepal, 1 from Nigeria, 1 from Pakistan, 1 from the Philippines, 1 from Senegal, 1 from Sierra Leone, 1 from South Africa, 2 from Shri Lanka, 2 from Tanzania, 1 from Uganda, and the rest of 12 genotypes were additionally added from the ICRISAT repository (Supplementary Table 1). Sowing (0.75 × 0.10 m spacing) of 176 accessions was completed in an alfa-lattice design with two replications at two locations (ICRISAT and GKVK). The sick plot for FW and SMD is situated at 13.08°N 77.58°E in Gandhi Krishi Vigyana Kendra (GKVK), Bengaluru, and at 17.54°N 78.28°E in ICRISAT, Hyderabad. In disease nursery, the susceptible check for SMD was ICP8863, and for FW it was ICP2376 at both the locations.

### 2.2 Phenotyping/ screening for FW and SMD

Phenotypic data for FW and SMD was recorded at the eight-leaf stage, pre-maturity stage, and maturity stage for 176-genotype panel in a rainy 2023-24. Additionally, we have used 143-genotype (from minicore collection) phenotypic data which was recorded in in rainy 2007-2008 and rainy 2008-2009 [5]. The disease score had been recorded using the disease incidence scale of 0-100%. The analysis was done by using the final compiled data (Supplementary Table 2). The disease incidence scale for both FW and SMD, ranging from 0% (highly resistant) to 100% (highly susceptible), was used to classify disease occurrence into four classes: 0-10% (resistant), 11-20% (moderately susceptible), 21-40% (susceptible), and >40% (highly susceptible). The overall disease incidence was calculated using the following formula [4]:

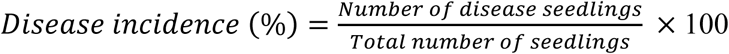

### 2.3 DNA isolation, library preparation and sequencing

DNA was extracted from all the genotypes using the NucleoSpin^®^ 96 Plant II (Macherey-Nagel) Kit, and DNA quality was checked on a 0.8% agarose gel quantity was estimated using Nanodrop spectrophotometer (Thermofisher Scientific, USA) as explained in [18], before the library preparation. Further, a single library was prepared for each sample using the TruSeq DNA Sample Prep kit. Genomic DNA was used to generate paired-end libraries for random sequencing on Illumina Hiseq 2000.

### 2.4 SNP filtration and quality control

Paired-end sequencing was performed for all samples using Illumina HiSeq 2000 instrument. Raw reads generated from these samples were subjected to quality filtering using FASTQC v0.12.0 [19], raspberry [20] and Trimmomatic v0.39 [21]. The low-quality reads (Phred score < 30; read length < 50 bases) and reads with adapter contamination were removed to generate a set of high-quality reads termed as clean data thereafter. High quality reads were mapped to the updated reference genome of pigeonpea (Cajca.Asha_v2.0) [13] using BWA ver. 0.5.9 [22] with default parameters. SNP calling was performed using GATK v.3.7 [23], as GATK has the best parameters for SNP calling. Biallelic SNPs with less than 20% missing calls, and cut-off of 5% on minor allelic frequency were proceeded for downstream analysis. Captured variants were annotated with SNPeff ver. 3.2 [24] and SnpSift ver. 3.2 [25].

### 2.5 Genetic diversity estimation

The diversity estimation for the 176 genotypes was done using the TASSEL software based on the nucleotide diversity. The neighbor-Joining statistics were considered to develop the diversity tree of 176 WGRS genotypes. Then the “NWK” extension file was used in the “iTOL” webtool (https://itol.embl.de/) to improve the graphics of the dendrogram.

### 2.6 Principal component analysis (Q matrix) and Kinship matrix (K matrix)

Principal component analysis (PCA) was conducted using the GAPIT package of RStudio [17]. The PCA was performed to identify the sub-populations within the selected panel of genotypes. The kinship matrix (K) depicts the genetic relationship in the selected population. K is used for correcting the false positives and most likely provides the true associations between marker and trait. K in GAPIT was calculated using the VanRaden Method [26]. The K was used as a random effect in some of the GWAS models [17].

### 2.7 Identification of marker-trait association

GWAS was performed in the GAPIT package of RStudio. The Bonferroni correction was implied in the models of GAPIT to confirm the statistically significant marker-trait associations (MTAs). The Bonferroni correction threshold was calculated using the following equation [27]:

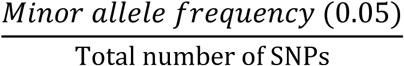

In this study, 5 models of GAPIT software were considered, viz., SUPER, BLINK, FarmCPU, CMLM, and MLM. All these models have different basic structures and the components, which contain fixed and random effects. PC score is used as a fixed effect for GWAS analysis. CMLM and SUPER generally contain higher statistical power than other models. MLM (mixed linear model), a single locus model, includes the kinship matrix (K) as a random effect along with the population structure (Q). This random effect is not included in the GLM model (general linear model). CMLM (compressed MLM), a single locus model, has enhanced statistical power than MLM because it makes the clusters of an individual into different groups, which used as an element to reduce the kinship matrix for random effect structure. FarmCPU (fixed and random model circulating probability unification) algorithms are used for multiple loci analysis. The FarmCPU model can efficiently manage both the false positives and false negatives. This model fixes both random and fix effects through an iterative approach; as this is a fix effect model, it has more computational efficiency than all other random models. In this model, associated markers are selected using the maximum likelihood method. BLINK (Bayesian information and Linkage-disequilibrium Iteratively Nested Keyway), a multi-locus model, is statistically superior to the FarmCPU model. BLINK uses the most significant marker as a reference and approximates the maximum likelihood using the Baysian information content (BIC). This model has more computational efficiency than the other models. In a SUPER, which is a single locus model, kinship (K) takes from the associated marker, which eliminates the confounding between K and some of the markers using the complementary kinship from the associated markers. SUPER uses the bin approach, while BLINK directly works on markers from LD (Linkage disequilibrium) to select the associate markers [17] [28].

### 2.8 Candidate gene identification and allele-mining of the putative markers

Based on MTAs identified, the nearest disease resistance related candidate genes were fetched. Due to the higher number of markers in the genotypic file, the Bonferroni correction threshold was higher in this study. Hence, to capture the minor putative candidate regions, the arbitrary threshold was also applied, and many minor MTAs were recovered between disease resistance-related candidate genes having significant allelic distribution. The allelic distribution of all the markers was checked in the whole population. The distribution of favourable and unfavourable alleles in the extreme genotypes was compared, and significant markers having associations with FW and SMD were selected.

The identified candidate genes from MTA region, were projected in FW and SMD disease resistance pathways.

### 2.9 Validation of the markers using KASP genotyping platform

For FW, total 30 markers from chromosome 11 were sent for the validation based on the identified allelic discrimination between FW resistant and FW susceptible genotypes. For SMD, Total 11 markers from chromosome 04 were validated based on the identified allelic distribution between SMD resistant and SMD susceptible genotypes. For the validation of FW markers, total 46 lines of ICPL85063 × ICPL87119 RIL population along with the parents were included in the validation panel.

For the validation of SMD markers, four BC_2_F_5_ and seven BC_2_F_6_ line of TS3R × ICPL20096 MABC population along with parents were included in the validation panel. All these lines are resistant to SMD (selected based on the phenotypic scores). Moreover, 94 reference set lines and 8 SMD resistant lines of ICPL11255 × ICP9174 F2:3 population were sent in the validation panel.

### 2.10 Identification of the haplotypes

The haplotype identification was done from the significant MTAs region. For the haplotype identification, we selected markers from the 1mb upstream and 1mb downstream region of significant MTAs to estimate the LD decay (r^2^). A subset of SNPs surrounding the identified significant MTAs was selected based on r^2^. The chosen SNPs account for the available LD variations surrounding the lead MTA. Wherever possible, these representative SNPs have diverse r^2^ values – low LD (r^2^: >0.3 and <=0.5), moderate LD (r^2^: >0.50 and <=0.70) and high LD (r2: >0.70). In case if no SNPs are satisfying the r^2^ threshold for a particular category, the other available SNPs were picked. However, only those SNPs that had homozygous calls were selected (removing heterozygous and missing calls). These representative SNPs were further used for clustering the haplotype groups, satisfying a basic criterion that all the unique patterns of these SNPs available in the genotypes GWAS panel are fully captured. Following this, ‘haplo-pheno’ analysis involved the comparison of FW and SMD means among all the haplotype groups that have at least 3 genotypes using one-way ANOVA coupled with Tukey’s HSD post-hoc test to identify the statistically different haplo-groups. The haplotype which exhibited highest levels of resistance were considered “resistant haplotypes”, while those displaying high levels of susceptibility were tagged as “susceptible haplotypes”.

## 3. Results

### 3.1 Phenotypic diversity for FW and SMD resistance

The phenotypic data for FW and SMD was recorded for 176-genotype panel in the rainy 2023-24 and for 143-genotype panel (From minicore collection) in rainy 2007-2008 and rainy 2008-2009 (Supplementary Table 2). Total 13 FW resistant lines with a mean of 6.4%, 12 moderately resistant lines with a mean of 16.8%, and 120 highly susceptible lines with a mean of 65.1% were obtained in the GKVK location. Total 7 FW resistant lines with a mean of 4.6%, 5 moderately resistant lines with a mean of 14.2%, and 155 highly susceptible lines with a mean of 72.6% were observed in the ICRISAT location.

Total 8 SMD resistant lines with a mean of 6.5%, 10 moderately resistant lines with a mean of 17.5%, and 119 highly susceptible lines with a mean of 55.9% were observed in the GKVK location. 34 SMD resistant lines with a mean of 3.8%, 23 moderately resistant lines with a mean of 15.9%, and 45 highly susceptible lines with a mean of 59.4% were observed in the ICRISAT location.

### 3.2 Sequencing statistics of genotypic data, genetic diversity and PCA

The sequencing data of 176 lines was aligned on the version 2 assembly of pigeonpea [13]. Maximum reads (10,061,8194bp) were obtained in ICP8211 genotype and minimum reads (24,159,362bp) were obtained in MAL-13 genotype. Total 69,195,482bp average reads were obtained. Maximum read mapped (99,964,766bp) were obtained in ICP8211 genotype and minimum read mapped (23,925,022bp) were obtained in MAL-13 genotype. Total 68,794,249bp average read mapped were obtained. Maximum percent mapped (99.44%) was obtained in ICP14722 genotype and minimum percent mapped (98.31%) was obtained in ICP11406 genotype. 99.27% average percent mapped was obtained. Maximum coverage of 16.2X was obtained in ICP8211 genotype and minimum coverage of 4.8X was obtained in MAL-13 genotype. Total 11.1X average coverage was obtained (Supplementary Table 3).

For the 176 lines, cluster I has 60 genotypes, cluster II has 42 genotypes, cluster III has 25 genotypes, cluster IV has 22 genotypes, and cluster IV has 27 genotypes. Similarly, in the PCA of the reference set, PC1 has 27 genotypes, PC2 has 41 genotypes, PC3 has 27 genotypes, PC4 has 56 genotypes, and PC5 has 25 genotypes. First two components (PC1 and PC2) account the maximum overall variability of 57.2% (Figure 1).

**Figure 1.**
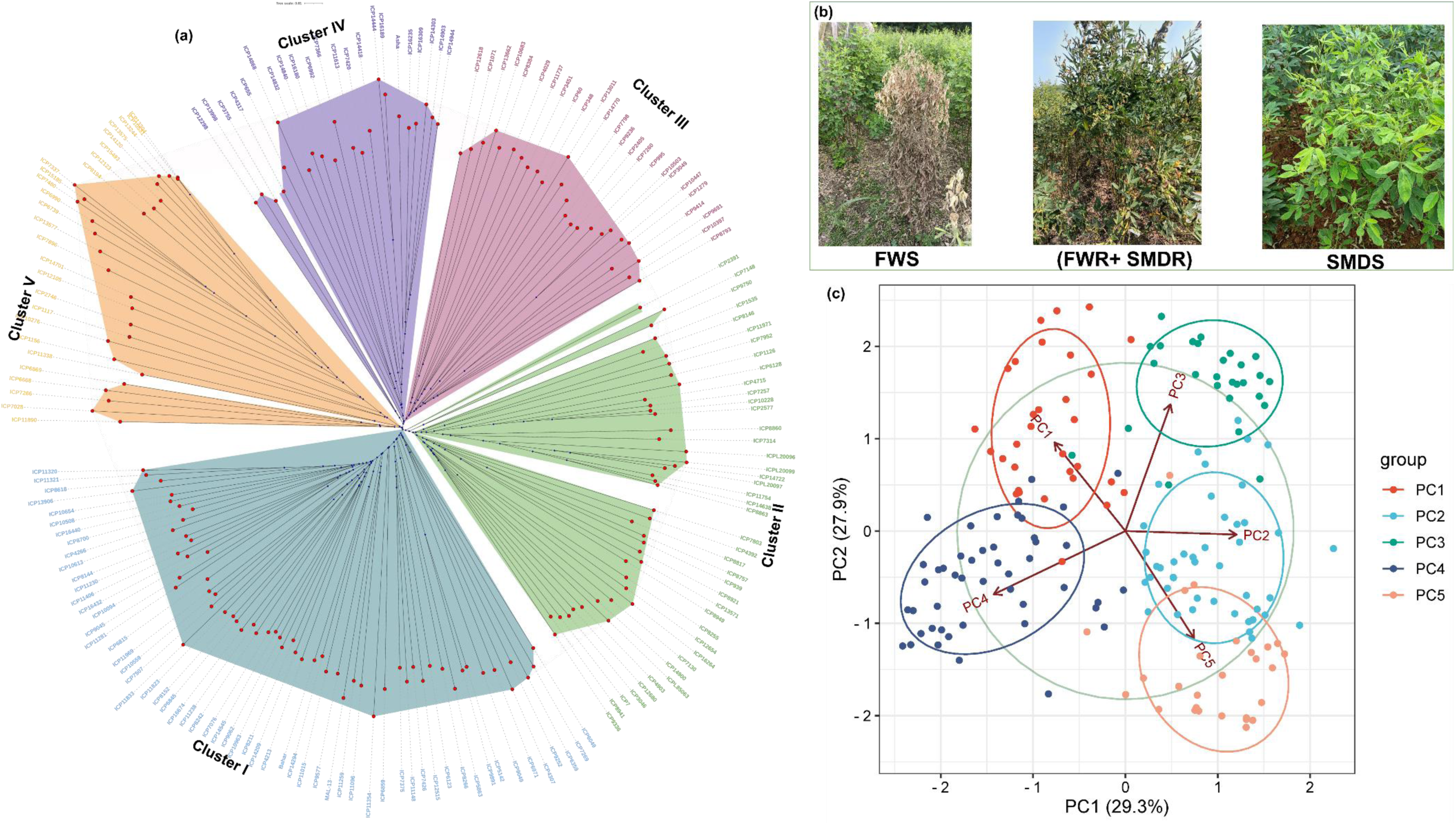
Representation of genetic divergence within the selected population. **(a)** Genetic diversity among 176-genotypes depicted using unweighted neighbour-joining tree method, this panel divided into five clusters, are represented in different colours. **(b)** Phenotype of FW susceptible, FW and SMD resistant, and SMD susceptible. **(c)** Sub-populations in different colours depicted as PCA plot, and first two components (PC1 and PC2) account the maximum overall variability of 57.2%

### 3.3 Identification of marker-trait association for FW and SMD

All the used models in 176-genotype panel, for both the locations (GKVK and ICRISAT) and pooled data, GWAS for FW observed a total of 5 putative markers above the Bonferroni correction threshold line. Similarly, for the 143-genotype panel, in all the used models, GWAS for FW observed a total of 23 MTAs which were above the Bonferroni correction threshold. Based on allelic distribution, we have selected 9 markers below the Bonferroni correction threshold. Moreover, GWAS was performed for the combined data of the 176-genotype and 143-genotype panel, and we have observed the total 13 putative markers above Bonferroni correction threshold line (Figure 2; Supplementary table 4).

**Figure 2.**
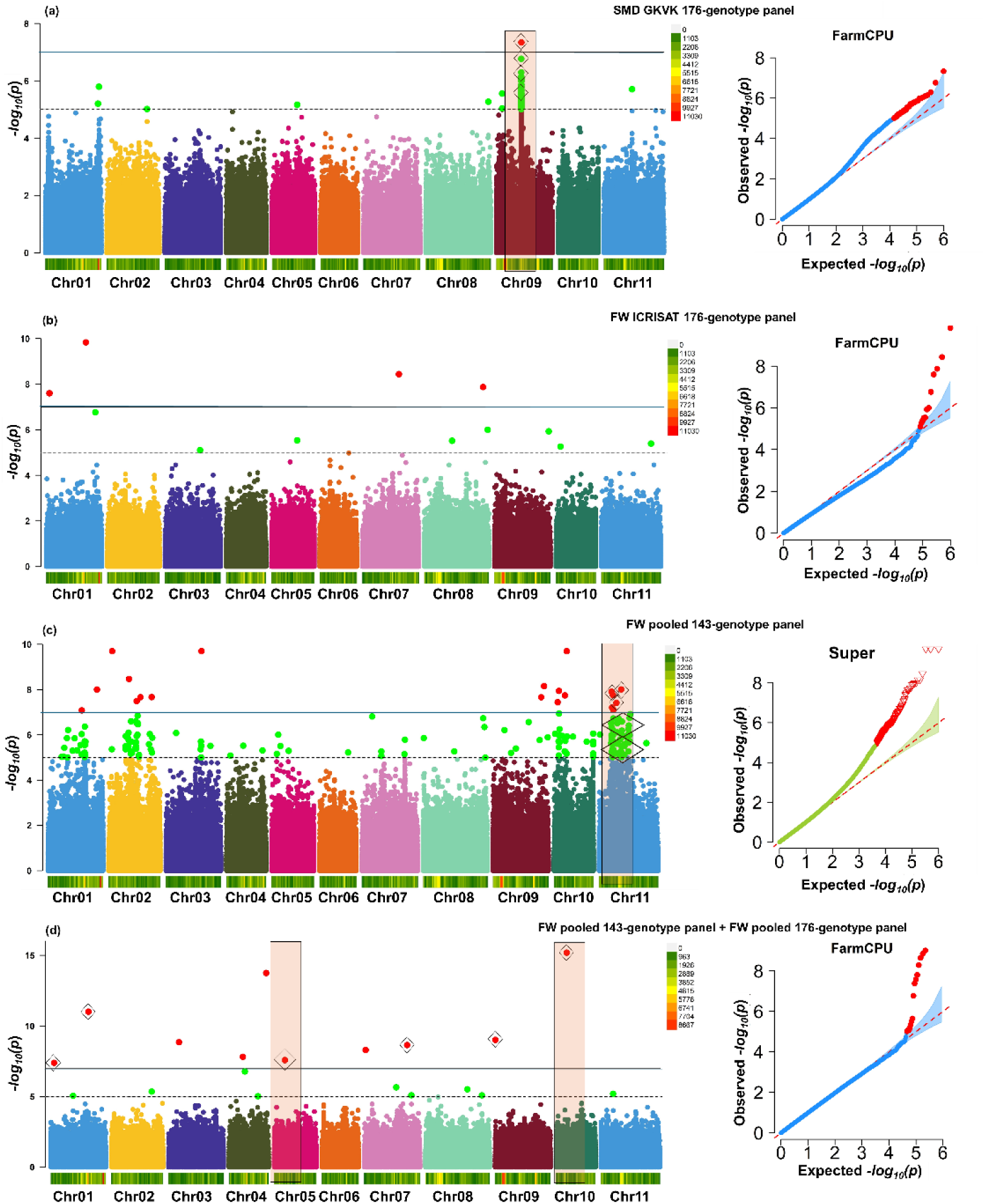
Identification of marker-trait associations for FW and SMD. Marker-trait associations (MTAs) for FW and SMD traits identified using genome-wide study in the 176-genotype and 143-genotype panel of pigeonpea: Manhattan plots and QQ plots showing association in these panel for FW and SMD. In Manhattan plots, solid black line represents Bonferroni correction line and red dots represents MTAs above this line, while dotted black line represents arbitrary threshold and the green dots represents MTAs above this line. The diamond box represents MTAs validated using phenotype-wise allelic distribution. The saffron colour highlighted boxes showing the highly significant regions. The x-axis of Manhattan plot represents chromosomes and SNP density plot and y-axis represents -log10(*p*) values. Upper right side of each Manhattan plots represents the scale to measure the SNP density of each chromosome. The x-axis of QQ plots represents expected -log10(*p*) values and y-axis represents observed -log10(*p*) values. in QQ plots, red dots represent all MTAs above arbitrary and Bonferroni correction threshold. **(a)** This Manhattan plot is showing SMD associated MTAs obtained from GKVK 176-genotype panel. 15 MTAs have shown the significant allelic distribution. Highly significant MTAs identified on chromosome 09 **(b)** This plot is showing the FW associated MTAs obtained from the ICRISAT 176-genotype panel data. **(c)** This Manhattan plot is showing the FW associated MTAs from 143-genotype panel. 43 MTAs have shown the significant allelic distribution **(d)** This Manhattan plot is showing the FW associated MTAs obtained from the combined analysis of 176-genotype panel and 143-genotype panel. Five MTAs have shown the significant allelic distribution.

From the region of FW significant markers, potential InDels were also identified. On chromosome 5, five significant InDels were identified in the promoter (4 InDels) and exonic (1 Indel) region of the *two component response regulator protein* (*Cc_11098*) gene. 13 resistant genotypes have shown favourable allele, while 32 susceptible genotypes have shown unfavourable allele (Figure 3). Among five InDels, four are housed in promoter region while one in exonic region. For *Cc05_10505987* marker in susceptible genotypes, the disruption has been observed (due to 1 bp deletion) in the transcription factor recognition site of *Dof-type zinc finger DNA-binding family protein* transcription factor (ATTAA). This transcription factor regulates the expression of pathogen recognition, signal transduction, and the production of defence compounds. This transcription factor controls the expression of the *two component response regulator gene* (*Cc_11098*) in the resistant genotypes.

**Figure 3.**
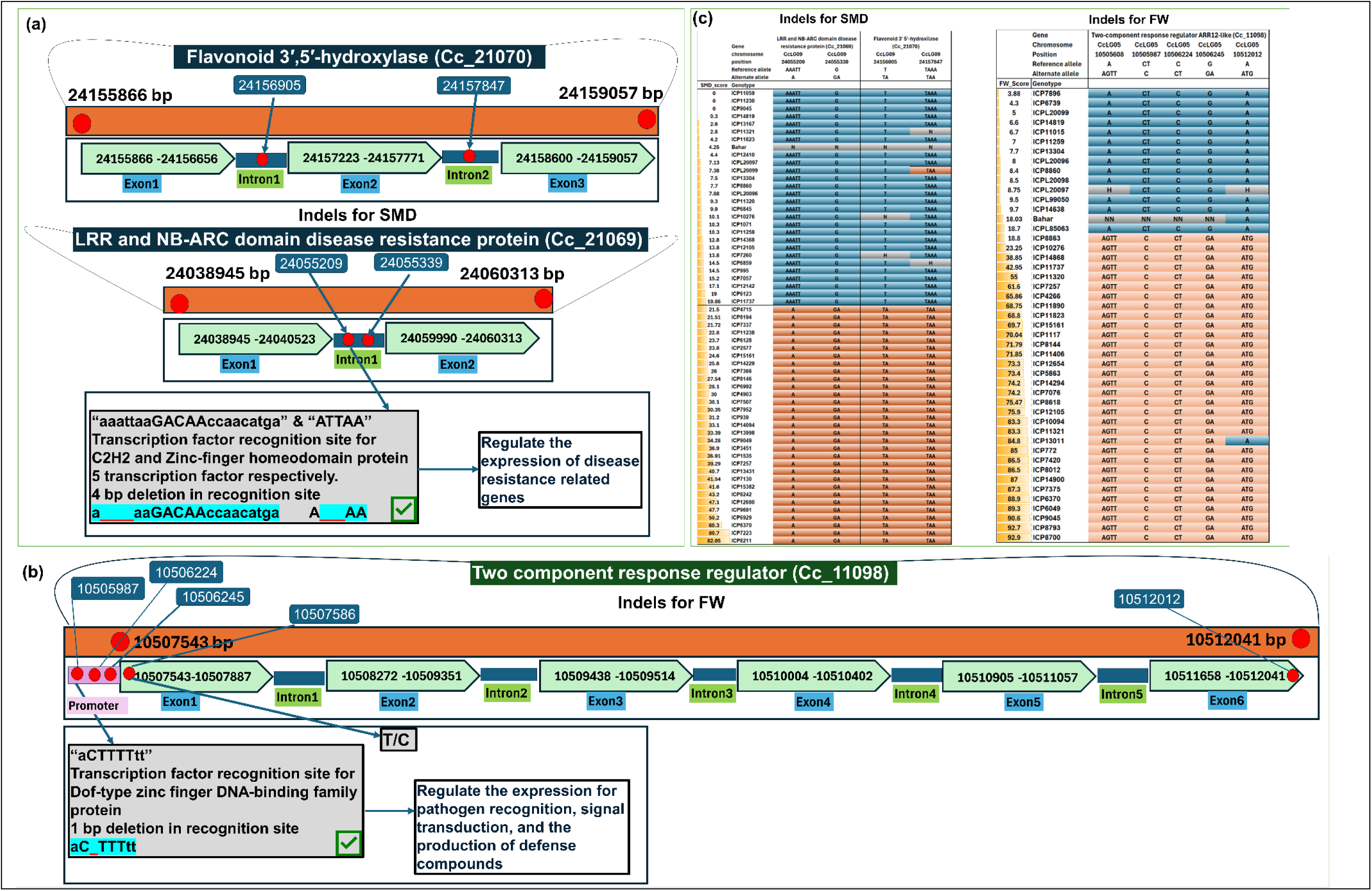
Identification of the InDels from FW and SMD associated region. The highlighted region with turquoise color is the transcription factor binding site and red underscore line indicated deleted region. **(a)** For FW, two InDels were identified in the intronic region of *Flavonoid 3’ 5’-htdroxylase* (*Cc_21070*) and two InDels were identified in the intronic region of *LRR and NB-ARC domain disease resistance protein* (*Cc_21069*). **(b)** For SMD, three InDels were identified in the promoter region and one indel identified in the exonic region f the *two-component response regulator* (*Cc_11098*)

From all used models in 176-genotype panel, for both the individual location data (GKVK and ICRISAT) and pooled data, GWAS for SMD observed a total of 3 putative markers above the Bonferroni correction threshold. Based on the allelic distribution and LD region, we have selected 14 additional markers below the Bonferroni correction threshold. For the 143-genotype panel, in all the used models, GWAS for SMD observed no putative markers above Bonferroni correction threshold. No putative markers were observed in the combined analysis of 176-genotype panel and 143-genotype panel. The detailed summary of these results is given in the Supplementary Table 4.

From the region of SMD significant markers, potential InDels were also identified. On chromosome 9, each two significant InDels were identified in the intronic region of *LRR and NB-ARC domain disease resistance gene* (*Cc_21069*), and *Flavonoid 3’ 5’-hydroxilase gene* (*Cc_21070*). 27 resistant genotypes have shown favourable allele, while 31 susceptible have shown unfavourable allele (Figure 3). For *Cc09_24055209* marker in susceptible genotypes, the disruption has been observed (due to 4 bp deletion) in the transcription factor recognition site of C2H2 (aaattaaGACAAccaacatga) and Zinc-finger homeodomain protein 5 transcription factor (ATTAA). Both the transcription factor regulates the expression of disease resistance related gene such as *LRR and NB-ARC domain disease resistance gene* (*Cc_21069*).

### 3.4 Identification of candidate genes involved in FW and SMD resistance mechanism

The candidate genes for FW markers and SMD markers were identified along with their function. The FW-associated markers were mostly found on chromosome 05, 10, and 11 (Table 1). The FW-associated markers are associated with disease-resistant related genes such *GATA transcription factor* (*Cc_26353*), *ubiquitin-conjugative protein* (*Cc_26039*), *receptor-like cytoplasmic kinase* (*Cc_26205*), and *serine-threonine protein kinase* (*Cc_26110*), *two component response regulator gene* (*Cc_11098*), *Dof-type zinc finger DNA-binding family protein* (*Cc_23564*) (Supplementary Table 4).

**Table 1.**
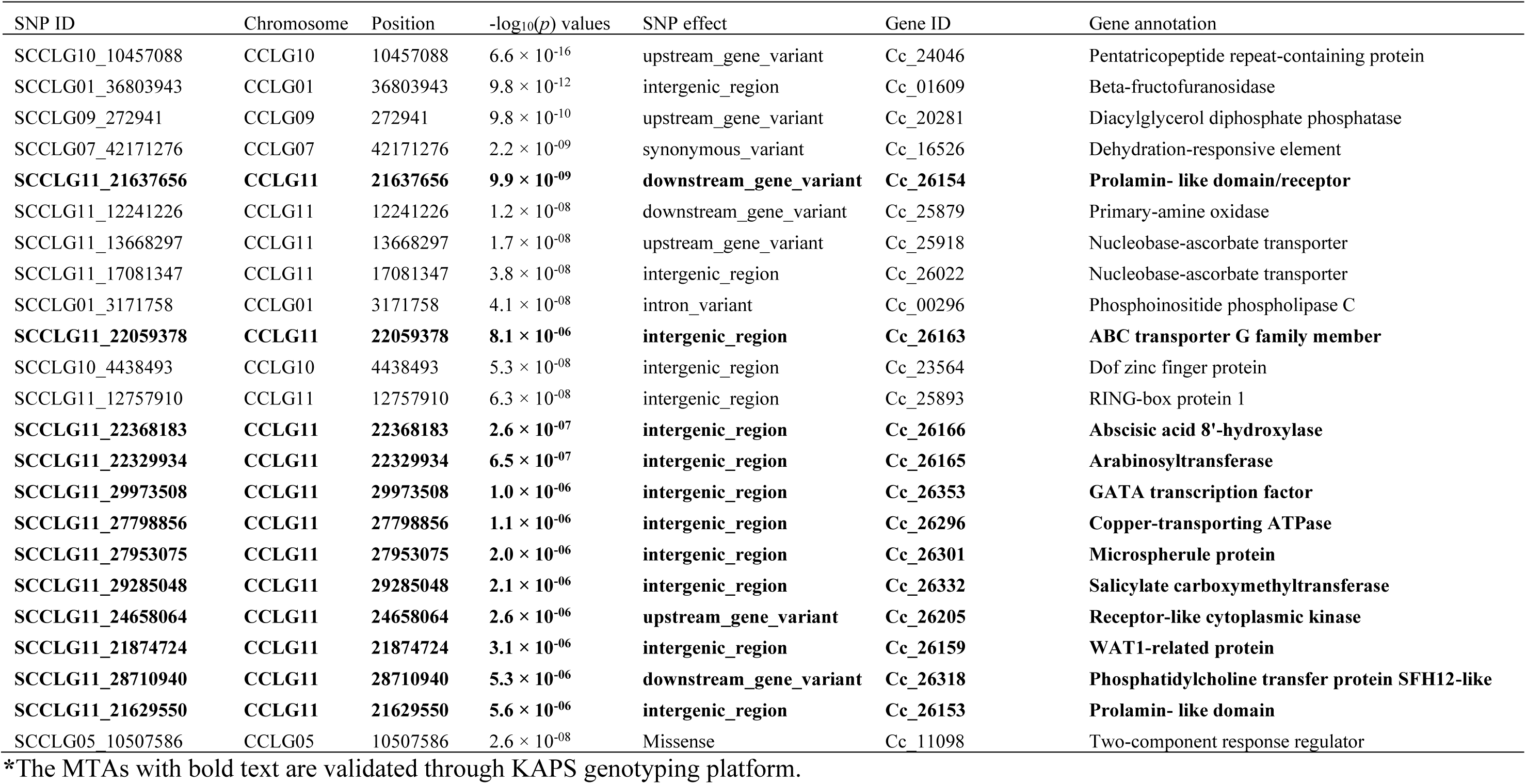
Identified significant MTAs for Fusarium wilt.

The SMD-associated markers are found on chromosome 09, and all the 3 markers are between three disease-resistant related genes, viz., *LRR and NB-ARC domain disease resistance protein* (*Cc_21069*) and *flavonoid 3’, 5’-hydroxylase* (*Cc_21070*) (Table 2). In the LD region of these markers, many copies of all three genes were found.

**Table 2.** Identified significant MTAs for sterility mosaic disease.

| SNP | Chromosome | Position | $-\log_{10}(p)$ values | Marker effect | Gene ID | Gene annotation |
| --- | --- | --- | --- | --- | --- | --- |
| **SCCLG09_24156069** | **CCLG09** | **24156069** | **$5.9 \times 10^{-6}$** | **synonymous_variant** | **Cc_21070** | **flavonoid 3', 5'-hydroxylase** |
| SCCLG09_24130097 | CCLG09 | 24130097 | $6.5 \times 10^{-6}$ | intergenic_region | Cc_21069 | LRR and NB-ARC domain disease resistance protein |
| SCCLG09_24147264 | CCLG09 | 24147264 | $6.7 \times 10^{-6}$ | intergenic_region | Cc_21069 | LRR and NB-ARC domain disease resistance protein |
| SCCLG09_24130102 | CCLG09 | 24130102 | $7.8 \times 10^{-6}$ | intergenic_region | Cc_21069 | LRR and NB-ARC domain disease resistance protein |
| SCCLG09_24162637 | CCLG09 | 24162637 | $7.9 \times 10^{-6}$ | upstream_gene_variant | Cc_21070 | flavonoid 3', 5'-hydroxylase |
| **SCCLG09_24131177** | **CCLG09** | **24131177** | **$4.6 \times 10^{-8}$** | **intergenic_region** | **Cc_21069** | **LRR and NB-ARC domain disease resistance protein** |
| **SCCLG09_24129144** | **CCLG09** | **24129144** | **$8.6 \times 10^{-6}$** | **intergenic_region** | **Cc_21069** | **LRR and NB-ARC domain disease resistance protein** |
| SCCLG09_24140843 | CCLG09 | 24140843 | $8.6 \times 10^{-6}$ | intergenic_region | Cc_21069 | LRR and NB-ARC domain disease resistance protein |
| SCCLG09_24194159 | CCLG09 | 24194159 | $9.1 \times 10^{-6}$ | intergenic_region | Cc_21071 | Putative disease resistance RPP13-like protein |
| **SCCLG09_24212766** | **CCLG09** | **24212766** | **$9.1 \times 10^{-6}$** | **intergenic_region** | **Cc_21071** | **Putative disease resistance RPP13-like protein** |

\*The MTAs with bold text are validated through KAPS genotyping platform.

For FW and SMD, we have projected disease resistance genes in plant-pathogen interaction pathways. In these pathways, 25 direct resistance related genes are found from our GWAS study. Some of these genes are; *GATA transcription factor 25*, *ABC transporter C family member 10*, *E3 ubiquitin-protein ligase PRT1*, *Receptor-like protein kinase HSL1*, *Serine-threonine kinase receptor-associated protein*, *LRR receptor-like serine/threonine-protein kinase*, *F-box/LRR-repeat protein 3*, *Plant intracellular Ras-group-related LRR protein 4*, *Actin-related protein 2/3 complex subunit 1B*, *Flavonoid 3’,5’-hydroxylase 1*, *Putative disease resistance protein*, *Putative disease resistance RPP13-like protein 1*, *Probable disease resistance protein*, *Putative serine/threonine-protein kinase-like protein CCR3*, *Potassium channel AKT1*, *Protein HIRA (Histone regulator protein)*, *Kinesin-like protein NACK1*, *Pentatricopeptide repeat-containing protein*, *Probable WRKY transcription factor 9*, *Mitogen-activated protein kinase*, *Autophagy-related protein 8C*, *WRKY transcription factor 6*, *NAC domain-containing protein 7*, and *Myb-related protein* (Figure 4).

**Figure 4.**
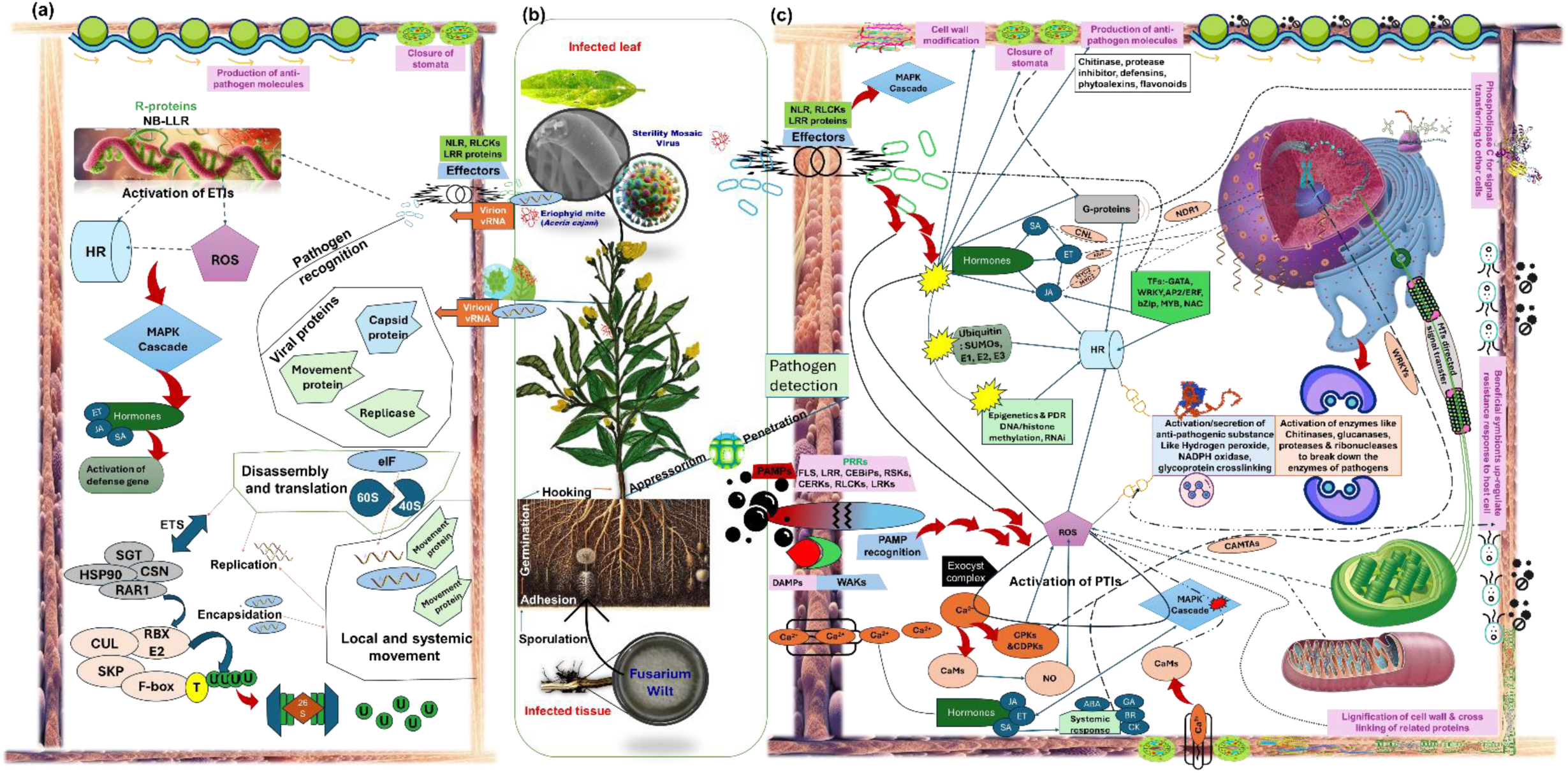
FW and SMD disease pathogen and plant interaction diagram. (**a)** Viral infection and resistance pathway. (**b)** The flow of infection for FW and SMD have shown. FW is a soil borne fungal pathogen mainly infect through the roots of plant and SMD is a viral pathogen, which transmit the plant through vector, eriophyid mite. (**c)** the plant cell showing the recognition mechanism of pathogens, primary signal transition mechanism, secondary signal transitions and activation of various resistant mechanisms. Black and blue lines showing the interaction between various components in resistance pathways. 25 direct resistance related genes are found in our GWAS study. Some of these genes are; GATA transcription factor 25, ABC transporter C family member 10, E3 ubiquitin-protein ligase PRT1, Receptor-like protein kinase HSL1, Serine-threonine kinase receptor-associated protein, LRR receptor-like serine/threonine-protein kinase, F-box/LRR-repeat protein 3, Plant intracellular Ras-group-related LRR protein 4, Actin-related protein 2/3 complex subunit 1B, Flavonoid 3’,5’-hydroxylase 1, Putative disease resistance protein, Putative disease resistance RPP13-like protein 1, Probable disease resistance protein, Putative serine/threonine-protein kinase-like protein CCR3, Potassium channel AKT1, Protein HIRA (Histone regulator protein), Kinesin-like protein NACK1, Pentatricopeptide repeat-containing protein, Probable WRKY transcription factor 9, Mitogen-activated protein kinase, Autophagy-related protein 8C, WRKY transcription factor 6, NAC domain-containing protein 7, Myb-related protein.

### 3.5 Allele-mining and identification of resistant and susceptible genotypes

For FW, the allele mining of all the markers was done in different sets of genotyping panels. The alleles of total 43 markers from the 143-genotype panel were successfully mined; out of these, 15 markers having favourable alleles are consistent in three genotypes and 28 markers having favourable alleles are consistent in four genotypes (Supplementary Figure 1). Henceforward, we have identified resistant genotype, viz., ICP14819, ICP11015, ICP13304, and ICP8860. The FW disease incidence scores of these genotypes are 6.6, 6.7, 7.7, and 8.4, respectively. There are 39 susceptible genotypes having unfavourable alleles (Supplementary Figure 1). Another set of alleles was mined from the combined analysis of a dataset having a 176- and 143-genotypes. In this dataset, we have identified five markers having favourable and unfavourable alleles. On chromosome 01, one marker was identified with favourable allele in ICP6739, ICP14819, ICP11015, ICP11259, and ICP85063 FW-resistant genotypes. On chromosome 07, one marker was identified with favourable allele in ICPL20099, ICP14819, ICP11015, and ICP8860 genotypes. On chr09 and chr10, one and two markers identified with favourable allele in ICPL20099, ICPL20096, ICP8860, ICPL85063, ICPL20097, and ICPL14838 were identified (Supplementary Figure 1). In addition, we have seen the LD region of these alleles contains many FW disease-resistance related genes. Some genotypes, such as ICPL20096, ICPL20099, and ICP8860 have multiple favourable regions, and hence, from these results, we can roughly conclude that the different resistant regions contribute to the various degrees of resistance in different genotypes in combinations.

For SMD, the allele mining was done in all the 17 markers. Among these, 15 markers on chromosome 09 having favourable allele combinations in 5 resistant and 10 moderately resistant genotypes. From this analysis, ICP8384, ICPL20096, ICPL20097, ICP13304, and ICP6815 were found to be SMD-resistant genotypes. Some genotypes like ICPL8255, ICP3046, ICP10508, etc. are moderately resistant. Interestingly, ICP13662, having most of these alleles in heterozygous allelic combinations, might be having a score of 50% because, SMD resistance is controlled by recessive genes (Supplementary Figure 1).

### 3.6 Marker validation using KASP genotyping platform

For FW, among 30 markers, 12 are validated on the ICPL85063 × ICPL87119 RIL population. Resistant parent ICPL87119 and 13 resistant inbred lines showed the favourable allele while susceptible parent ICPL85063 along with 9 susceptible inbred lines showed the unfavourable alleles in 12 KASP markers, viz., snpCC00128 (*Cc11_21629550*), snpCC00129 (*Cc11_21637656*), snpCC00130 (*Cc11_21874724*), snpCC00132 (*Cc11_22059378*), snpCC00133 (*Cc11_22329934*), snpCC00134 (*Cc11_22368183*), snpCC00135 (*Cc11_24658064*), snpCC00138 (*Cc11_27798856*), snpCC00139 (*Cc11_27953075*), snpCC00140 (*Cc11_28710940*), snpCC00144 (*Cc11_29285048*), and snpCC00146 (*Cc11_29973508*) (Figure 5).

**Figure 5.**
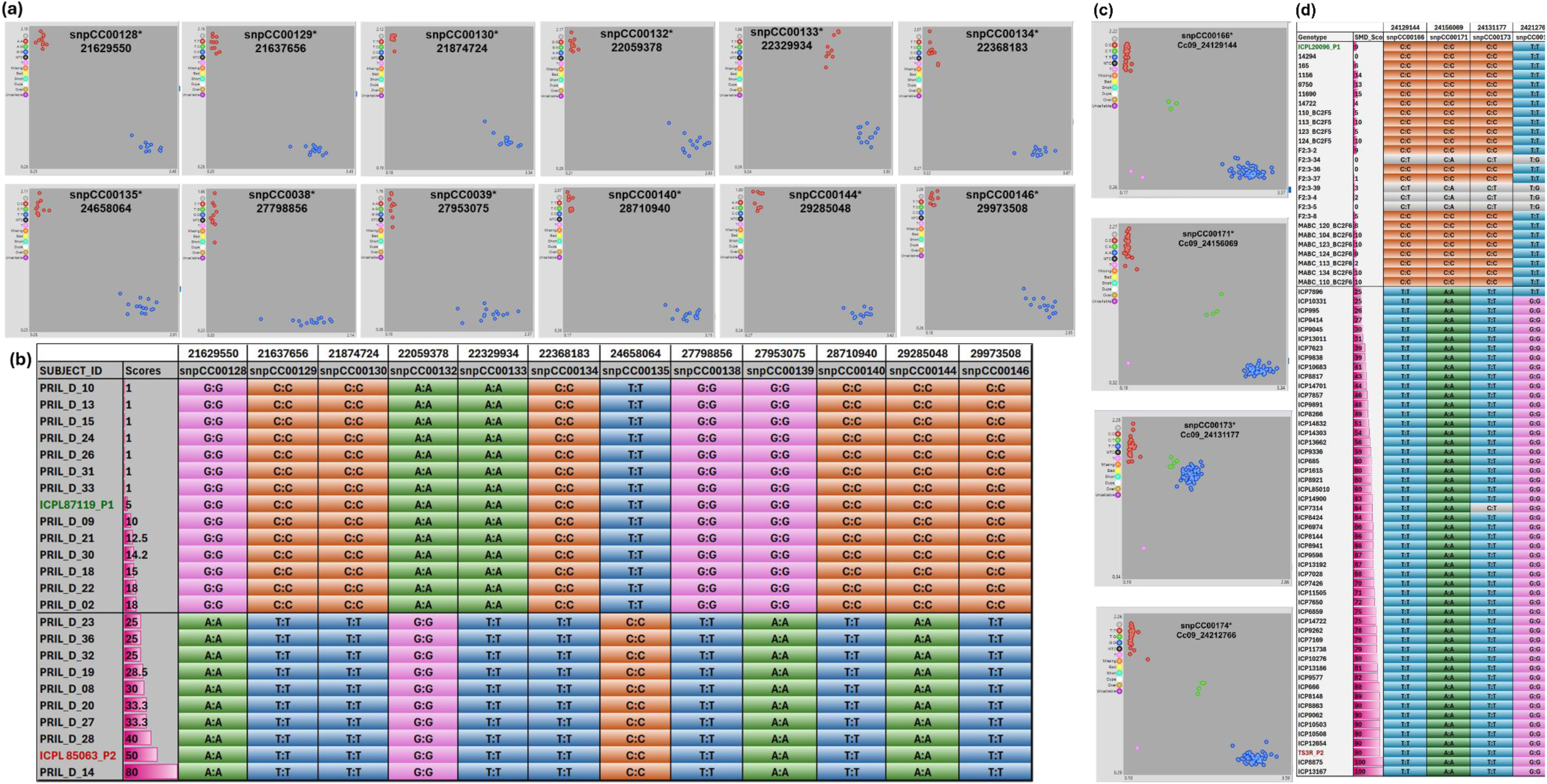
The pictorial representation of the marker validation using KASP genotyping assay. **(a)** For FW, the validation conducted on a RIL population (ICPL 85063 × ICPL87119) containing 13 resistant lines and 9 susceptible lines including parents. Red color indicates the unfavourable allele, blue color indicates the favourable and green color indicates the heterozygotes alleles. Star indicates Intertek IDs **(b)** 12 markers validated from the intergenic region nearby to the genes, namely., *Prolamin-like domain* (*Cc_26154*), *ABC transporter G family member* (*Cc_26163*), *Abscisic acid 8’-hydroxylase* (*Cc_26166*), *Arabinosyltransferase* (*Cc_26165*), *GATA transcription factor* (*Cc_26353*), *Copper-transporting ATPase* (*Cc_26296*), *Microspherule protein* (*Cc_26301*), *Salicylate carboxymethyltransferase* (*Cc_26332*), *Receptor-like cytoplasmic kinase* (*Cc_26205*), *WAT1-related protein* (*Cc_26159*), *phosphatidylcholine transfer protein SFH12-like* (*Cc_26318*). **(c)** For SMD, the validation conducted on a MABC population (TS3R × ICPL20096) of 2 generations lines (4 from BC_2_F_5_ and 7 from BC_2_F_6_) including parents and reference set genotypes. Red color indicates the favourable allele; blue color indicates the unfavourable and green color indicates the heterozygotes. Star indicates Intertek IDs **(d)** Four markers validated from the intergenic region between three disease resistance genes, Viz., *LRR and NB-ARC domain disease resistance protein* (*Cc_21069*), *Flavonoid 3’, 5’-hydroxylase* (*Cc_21070*), and *Probable disease resistance protein* (*Cc_21071*)

Similarly, for SMD, among 11 KASP markers, 4 markers were successfully validated, viz., snpCC00166 (*Cc09_24129144*), snpCC00171 (*Cc09_24156069*), snpCC00173 (*Cc09_24131177*), snpCC00174 (*Cc09_24212766*) (Figure 5). SMD resistant parent ICPL20096 along with the four BC_2_F_5_ and seven BC_2_F_6_ line of TS3R × ICPL20096 MABC population showed the favorable allele. Six SMD resistant lines of pigeonpea reference set and 8 SMD resistant lines of ICPL11255 × ICP9174 F2:3 population showed the favourable alleles. While susceptible parent TS3R, and 51 susceptible lines of pigeonpea reference set showed the unfavourable alleles (Figure 5).

### 3.7 Identification of resistant haplotypes

For FW, based on the indel identification, high PVE%, and marker validation, three important MTAs were selected, viz., *Cc05_10507586* from chromosome 5, *Cc10_4438493* from chromosome 10, and *Cc11_21637656* from chromosome 11 respectively. For SMD, based on marker validation, indel identification and high PVE%, *Cc9_24131177* from chromosome 09 was selected for the haplotype identification. Based on the r^2^ values, representative markers were selected (Supplementary Table 5). For the *Cc05_10507586* region of chromosome 05, H1 (TCATAATCGAT) is identified in 37 genotypes, H2 (CTGCGGCTACA) is identified in 11 genotypes, H3 (TCATAATCGCA) in 7 genotypes, H5 (CCATAATCGCA) in 4 genotypes, and H6 (TCGCGGCTGAT) in 8 genotypes. All the haplotypes from this region are identified in most of the FW susceptible genotypes. From the region *Cc10_4438493* of chromosome 10, H1 (CAC) is identified in 128 genotypes, H2 (CAT) in 16 genotypes, H3 (TAC) in 5 genotypes, H4 (TGT) in 3 genotypes, H5 (TAT) in 7 genotypes, and H6 (CGT) in 3 genotypes. Among these 5 haplotypes, H4 is only identified in resistant genotypes, and the other 4 haplotypes are mostly in susceptible genotypes. Similarly, for *Cc11_21637656* of chromosome 11, H1 (TCGCCC) is identified in 143 genotypes, H3 (CTTTTT) in 3 genotypes, H4 (CCGCCC) in 7 genotypes, and H6 (TCTTTC) in 2 genotypes. Among these 4 haplotypes, H3 is identified only in resistant genotypes, and the other 3 haplotypes are mostly in susceptible genotypes (Figure 6). In conclusion, H4 from *Cc10_4438493* region and H3 from *Cc11_21637656* region can be called “resistant haplotype”. H4 from *Cc10_4438493* is identified in four FW resistant genotypes viz., ICPL20096, ICP14638, ICP14722, and ICP8863. H3 from *Cc11_21637656* is identified in three FW resistant genotypes viz., ICP11015, ICP13304, and 14819 (Supplementary Table 6).

**Figure 6:**
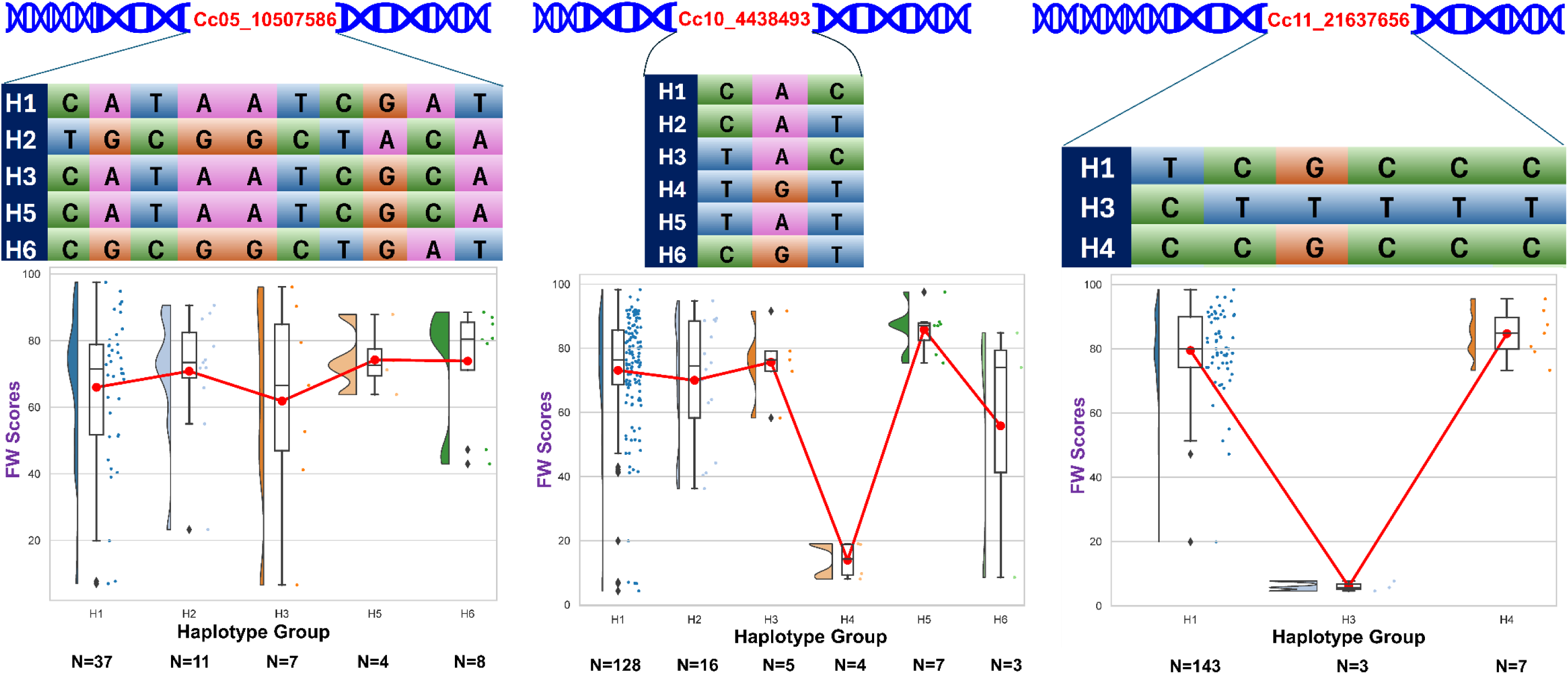
Identification of haplotypes from the significant MTA regions for FW. From *Cc05_10507586* region of chromosome 5, five haplotypes were identified. All five were identified in susceptible genotypes. From region *Cc10_4438493* in chromosome 10, one haplotype (H4) was identified only in resistant genotypes. Similarly, From A region *Cc11_21637656* of chromosome 11, six haplotypes were identified. Among six haplotype, one haplotype (H3) was identified only in resistance genotypes. N indicates number of genotypes having respective haplotypes. Red line indicates mean value connection value. Line inside the box indicated statistical mean (Outliers excluded).

For SMD, four SNPs viz., snpCC00166 (*Cc09_24129144*), snpCC00171 (*Cc09_24156069*), snpCC00173 (*Cc09_24131177*), snpCC00174 (*Cc09_24212766*) were considered for haplotype identification. The haplotype “H1” (CCCT) was identified in resistant lines while the haplotype “H2” (TATG) was identified in susceptible lines. Therefore, H1 (CCCT) is called a resistant haplotype while H2 (TATG) can be called as a susceptible haplotype (Figure 5). Both the haplotypes have the favorable (H1) and unfavorable (H2) calls from the same set of markers.

## 4 Discussion

The pigeonpea crop has an ideal nutritional content, can greatly tolerate environmental stresses, and has high biomass, which also helps to restore and recycle the important nutrients in soil. Pigeonpea has huge potential to improve the quality and quantity of foodgrains in India as well as other parts of the world, especially African countries [29]. However, few yield-reducing traits need to be strengthened. FW and SMD are the most significant yield-reducing barrier to pigeonpea. Due to the persistent and evolving feature of Fusarium udum Butler (A pathogen of Fusarium wilt), the identification of multiple resistance regions is highly necessary [30]. SMD is an endemic to the Indian subcontinent due to continuous enhancement in the infected area and yield losses [31]. Each disease resistant genomic region contains a unique and combined set of disease-resistance related genes. All the disease resistance related regions may not be present in single genotypes. Therefore, significant resistance regions from different sources need to be identified and combined in the genetic background of elite cultivars. The use of chemical formulations to control the disease incidence has been spoiling the soil and environmental health. The use of conventional breeding techniques take time to develop a disease-resistance variety. Moreover, the continuously evolving nature of pathogens leads to resistance breakage. GWAS investigates whole genomic variation, and it can identify multiple disease resistance regions such as candidate genes and linked markers [32]. The identified regions from different GWAS studies can be combined in the elite cultivars through marker-assisted selection and marker-assisted backcross selection approaches, which will reduce the cost and breeding time to develop multi-race resistance cultivars. One of the association studies of wheat revealed that the known resistant genes in most of the varieties are no more effective against the continuously evolving leaf rust pathogen [33]. Further, they have identified novel genes proving complete resistance in three wheat varieties. Moreover, they concluded that some other unknown resistance region provides resistance to the local cultivars of wheat. In our study also, we have identified similar results.

In this study, we identified genomic regions associated with FW and SMD resistance using a 176-genotype panel and 143-genotype panel. We have used the updated assembly of the reference genome ICPL87119 (Asha) for SNP calling in sequenced panel leading to 10,45,509 SNP variants after data trimming and filtering. After applying a 0.05 site minimum allele frequency and a 0.25 maximum heterozygous proportion filter, a total of 869447 SNPs were used for the actual association study for both panels.

Total 49 MTAs were identified for FW from WGRS genotyping data of the 176-genotype panel, 143-genotype panel, and combined analysis of both the panels were obtained. A cluster of markers on chromosome 11 follow allelic combination trend in three important FW resistant genotypes, namely ICP14819, ICP11015, and ICP13304. This result shows that the identified region on chromosome 11, is the source of resistance for these three genotypes. The combined phenotypic variation explained (PVE) is 30% for this region. For FW, Alleles from chromosomes 01, 07, 09, and 10, were also mined. Many diseases-resistance-related genes were also recovered from these regions. The additional markers below Bonferroni correction threshold were identified in the association study of humans to identify the total circulating tau level (used as an endophenotype to identify genetic risk factors related to neurological disease in humans) [34]. In this study, researchers detect the 14 novel loci associated with total circulating tau at p< 5 × 10^-7^ (from the below Bonferroni correction threshold). Similarly, in our study, we have identified addition FW-associated regions from the below Bonferroni correction threshold.

Candidate genes are the major findings of GWAS studies. These candidate genes are responsible for driving the disease resistance pathways from identifying pathogens to digesting them completely. Moreover, many genes used to keep these stress memories. Identified FW disease resistance genes from the mined region are: *GATA transcription factor* (*Cc_25907*), *E3 ubiquitin-protein ligase* (*Cc_07896*), *receptor-like cytoplasmic kinase* (*Cc_26205*), *WRKY transcription factor* (*Cc_26159*), *serine-threonine protein kinase* (*Cc_26110*), and *two component response regulator gene* (*Cc_11098*).

The crosstalk network analysis in rice revealed that the *GATA transcription factor* co-expressed with the *WRKY* and *NAC transcription factors* to regulate the plant disease mechanism [35]. In Arabidopsis, the contribution of *GATA family proteins* was recorded in hypersensitive response against pathogen *Pseudomonas syringae* [36]. Multidrug resistant protein in plants controls the stomatal transpiration and helps in detoxification of plants from unwanted materials [37]. The overexpression and translocation of berberine (a multidrug resistant chemical) to the rhizome was recorded in *coctus japonica* to perform the antimicrobial activities [38]. *Ubiquitin-conjugative protein* (E2) is the conjugative protein is triggered after the activation of E1 enzyme followed by the ligation of *E3 enzyme* in the ubiquitin proteosome system [39]. The expression of rice *E2 gene* was observed after the induction of N-acetylchitooligosaccharide (elicitor) in suspension cultured rice cells [40]. N-acetylchitooligosaccharide is a component of chitin released from the cell wall of a fungal pathogen during the infection in plant cells [41]. Due to the strong role of *receptor-like cytoplasmic kinase* (RLCKs) in PAMP-triggered immunity (PTI) and effector-triggered immunity (ETI), pathogens try to manipulate them to restrict the defence response [42]. RLCKs activate downstream signal and regulators such as hormones, Ca^2+^ signalling, accumulation of reactive oxygen species (ROS), activation of *mitogen-activated protein kinases* (MAPKs), and activation transcription factor (Figure 4). Multiple genes from the RLCKs family were knocked. These results showed the role of RLCK family protein against the oomycete pathogen of *nicotina benthamiana* [43]. Moreover, the molecular evidence for the activation of downstream defence-related components. *Serine-threonine protein kinase* works as an extracellular receptor to sense the presence of pathogen. In Arabidopsis, lines containing T-DNA of mutant *PBL13* (*Serine-threonine protein kinase*) genes were found to have higher expression of this variant than the normal one. This suggests the negative regulation of this gene in Arabidopsis [44]. The diagrammatic representation of the plant-pathogen interaction is shown in Figure 4.

On chromosome 5, we have identified a *Two component response regulator gene* (*Cc_11098*) associated with significant MTA *Cc_10507586*. This gene is involved in the signalling of external stimuli, such as pathogen attack, and passing these signals to activate the downstream defence mechanism. Two component system contains histidine protein kinase, which senses the external signal, and a response regulator, which transfers these signals to active defence response machinery [45]. Moreover, we have identified five InDels in *Two component response regulator gene* (*Cc_11098*). We have identified 1 bp deletion (at 10505987 bp position of chromosome 05) in the promoter region of this gene containing a transcription factor binding site (aCTTTTtt) for *Dof-type zinc finger DNA-binding family protein* (*Cc_23564*). In our study we have identified the region on chromosome 10 containing *Dof-type zinc finger DNA-binding family protein* (*Cc_23564*) with 57% of phenotypic variation explained (PVE). This transcription factor may be involved in the activation of *Two component response regulator gene* (*Cc_11098*).

In the view of above, we suggest that the regions from chromosome 5 (*Cc_10505987*) and chromosome 10 (*Cc_4438493*) are strongly associated with the FW resistance. Moreover, the favourable allele from chromosome 10 (*Cc_4438493*) is not present in FW resistant donor ICPL87119, however it is present in another FW donor, ICPL 20096. The favourable region *Cc_21637656* is present in ICPL87119 but not in ICPL20096. The favourable indel allele from chromosome 5 (*Cc_10505987*) is present in ICPL87119 as well as ICPL 20096. Therefore, we proposed the allele pyramiding using both the FW donors to develop highly resistant FW line (Figure 7).

**Figure 7.**
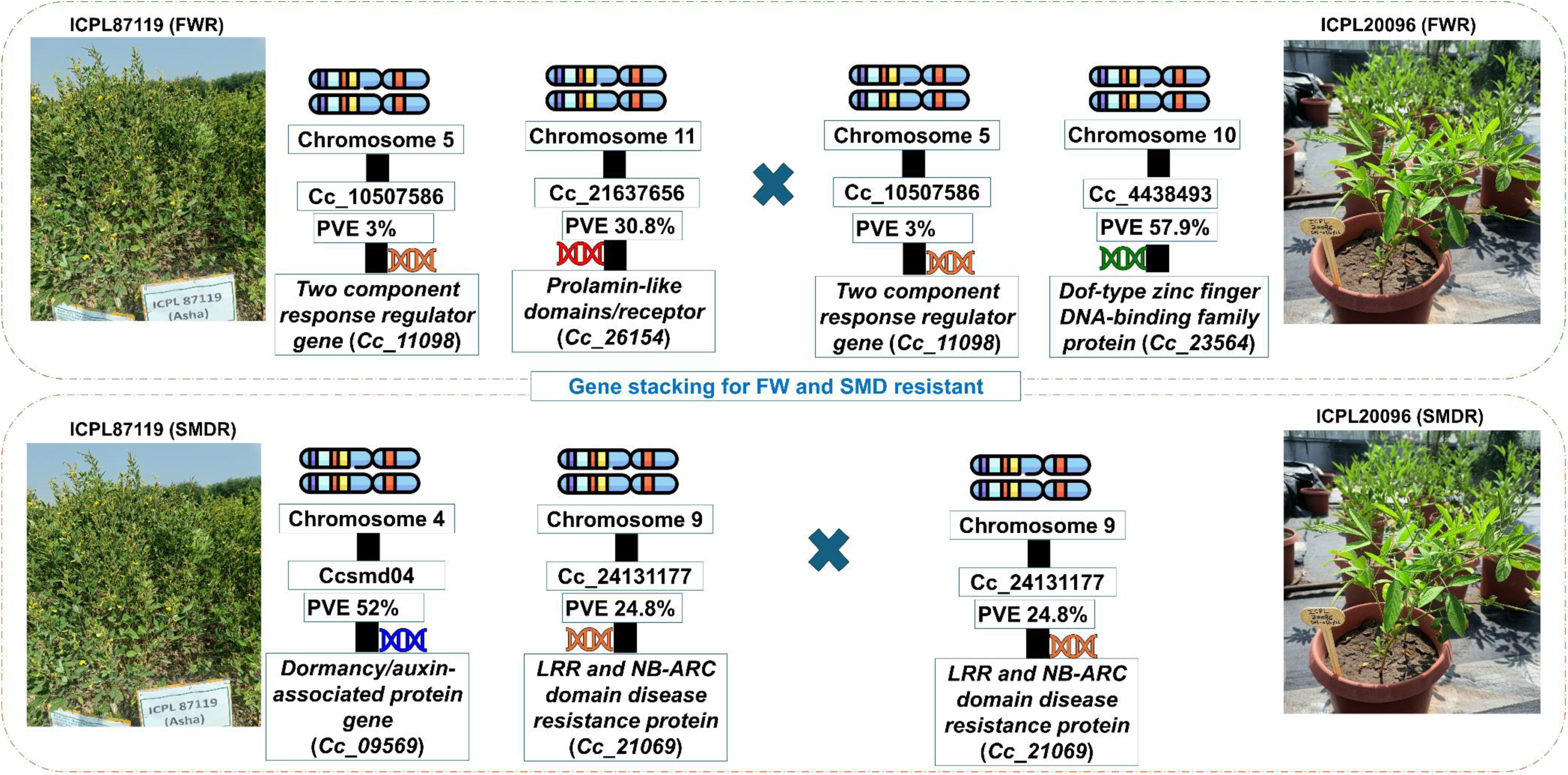
Proposed strategy for gene stacking for FW and SMD resistant. For FW, *Cc_10507586* region from chromosome 5 is identified in ICPL87119 and ICPL20096 donors. A region *Cc_21637656* from chromosome 11 is identified in the ICPL87119 but not in the ICPL20096. Similarly, a region *Cc_4438493* from chromosome 10 is identified in the ICPL20096 but not in the ICPL87119. For SMD, *Cc_24131177* region is identified in both the SMD resistance donors (ICPL20096 and ICPL87119). *Ccsmd04* region from chromosome 4 has identified in ICPL87119 but not in the ICPL20096.

A total 17 SMD resistance-related regions were obtained from WGRS genotyping data of the 176-genotype panel, 143-genotype panel, and combined analysis of both the panels. Among these 15 of them showed significant allelic distribution in resistant and susceptible genotypes, as shown in Supplementary Figure 2. A cluster of markers on chromosome 09 follow some allelic combination trend in 6 important SMD resistant genotypes, viz., ICP8384, ICPL20096, ICPL20097, ICPL87119, ICP13304, ICP6815, and ICP8860. The result shows that the region from chromosome 9 is the source of SMD resistance for these genotypes. The combined phenotypic variation explained (PVE) is 30% for this region. Many diseases-resistance-related genes were also recovered from these regions.

Three most important identified candidate genes are: *LRR* and *NB-ARC domain disease resistance protein* (*Cc_21069*), *flavonoid 3’, 5’-hydroxylase* (*Cc_21070*), and *putative disease resistance RPP13-like protein* (*Cc_21071*). *LRR* and *NB-ARC domain disease resistance protein* are directly and indirectly regulating plant disease resistance. In direct mode, this protein interacts with pathogen-derived molecules, while in indirect mode, effector molecules of pathogens are identified by this protein [46]. The nucleotide binding site of *NB-ARC domain* regulates the activity of R proteins [47]. The *Cysteine-rich-receptor-like secreted protein* (CRRSP) and *NB-ARC* complex triggered the oxidative state, cell death process, and associated resistance responses observed in rice [48]. *Flavonoid 3’, 5’-hydroxylase* is a key enzyme to control the hydroxylation pattern in flavonoid biosynthesis. The over-accumulation of a flavonoid in the cell of the rice line was observed through scanning electron microscopy and spectrophotometry after the infection of the Bacterial leaf blight pathogen *Xanthomonas oryzae pv. oryzae* [49]. *Putative disease resistance RPP13-like protein* is highly important as disease resistant gene. In Arabidopsis, this gene was found to be independently responsible for resistance to pathogens. No accumulation of salicylic acid is required to activate the disease resistance response [50]. Moreover, they found that the *RPP13 gene* restricted the infection of five isolates of *Peronospora parasitica* (At) in a transgenic Arabidopsis. In one study of durian (a tropical fruit crop), a resistant gene analogue (RGA) was analysed for functional characterisation of resistant genes [51]. Among the well-known disease resistance protein, the highest copies (207) of *putative disease resistance RPP13-like protein* were found in this RGA analysis.

On chromosome 9, we have identified *LRR* and *NB-ARC domain disease resistance protein* (*Cc_21069*), *flavonoid 3’, 5’-hydroxylase* (*Cc_21070*), and *putative disease resistance RPP13-like protein* (*Cc_21071*) gene associated with significant MTAs Cc_24131177. We have identified two InDels in the intronic region of *LRR* and *NB-ARC domain disease resistance protein* (*Cc_21069*), two InDels in *flavonoid 3’, 5’-hydroxylase* (*Cc_21070*). In *LRR* and *NB-ARC domain disease resistance protein* (*Cc_21069*) gene, indel Cc_ 24055209 has shown 4 bp deletion. Due to this 4 bp deletion, disturbance in the transcription factor binding site for C2H2 zinc finger transcription factors (aaattaaGACAAccaacatga) and Zinc-finger homeodomain protein 5 (ATTAA) transcription factor have identified. C2H2 zinc finger transcription factor (TFs) and Zinc-finger homeodomain protein 5 play an important role in plant disease resistance by directly regulating the expression of pathogenesis-related genes [52–53]. Although, these transcription factor binding sites are in intronic region, but some of the studies report that the presence of the transcription factor recognition site in the intronic region of the gene could play an important role in the gene expression regulation by influencing splicing mechanisms and potentially modifying the rate of transcription elongation within the body of gene. These sites can act as regulatory elements within the intron to control the transcription of the gene and their processing into mature mRNA [54]. Few earlier studies have confirmed that the conserved transcription factor binding sites are required for the appropriate expression of the genes in *Caenorhabditis elegance* [55–58]. In *Arabidopsis thaliana*, the second intron of AGAMOUS (AG) gene contains cis-acting DNA sequences that bind the transcription factors LEAFY and WUSCHEL which control floral fate and floral-specific expression of AG, respectively [59]. Such types of introns that act as a co and post-transcriptional regulation are called as intron-mediated enhancement (IME). Earlier, this phenomenon was reported in maize genes *Alcohol dehydrogenase1* and *Bronze1* [60], and *Shrunken1* [61]. In *Arabidopsis thaliana* it was reported in gene *Phosphoribosylanthranilate transferase1* [62].

In this GWAS study, we have identified a highly promising region (*Cc_24131177*) from chromosome 9 in SMD resistant donor ICPL 20096 and ICPL 87119. To identify SMD resistant regions, we have also performed QTL-seq analysis using ICP 8863 × ICPL 87119 RIL population. In our QTL-seq study we have reported a strongly associated SMD resistant region on chromosome 4. We checked this region in our other SMD-resistant donor ICPL20096 using KASP genotyping, and we found that this region is lacking in this donor [63]. Therefore, we strongly recommend for gene pyramiding by crossing ICPL 87119 and ICPL 20096 SMD resistant donors to develop a highly SMD resistant lines. Moreover, as we have identified donor regions for FW in the same donors, we recommend conducting gene stacking to develop the highly resistant FW and SMD lines (Figure 7).

ICP14819, ICP11015, ICP13304, ICP8860, ICP6739, ICP11259, ICPL85063, ICPL20099, ICPL20096, ICPL20097, ICPL87119, and ICP14638 FW resistant genotypes were obtained in this study. Similarly, ICP8384, ICPL20096, ICPL20097, ICP13304, ICP6815, ICPL87119 and ICP8860 SMD-resistant genotypes were identified from this study. Among these genotypes, ICPL20096, ICPL20097, and ICP13304 were found to be resistant for both FW and SMD. Sequencing based-bulked sergregant analysis (BSA) was performed with the 20096 × ICPL332 RIL population, where ICPL20096 was used as a resistant parent for both FW and SMD [14]. Additionally, to identify the nonsynonymous region, ICP20097 was one of the resistant parents for both the diseases. Again, in the genotyping-by-sequencing (GBS) based QTL mapping study, ICPL20096 and ICPL20097 were used as resistant parents for SMD [16]. In another GBS-based QTL mapping study, ICPL20096 and ICPL85063 were used as a resistant parent for FW [64]. For the validation of FW and SMD diagnostic kit markers, one BC_1_F_1_ population was developed by crossing ICPL20096 FW and SMD resistant parent with BDN711 [65]. Additionally, ICPL20099 as a FW and SMD resistant parent was used for the validation of FW and SMD markers. The evaluation of minicore collection for FW and SMD was carried out at the sick plot of Dharwad, India [66]. From this screening, ICP8860 was found resistant for SMD and FW, and ICP13304 was found resistant for FW only.

One QTL from the RIL population (ICPB2049 × ICPL99050) for FW was identified in all previous studies on chromosome 11, having a marker interval 16.8 Mb to 20.6 Mb with 10.04% PVE and 3.25 LOD score [64]. The identified region on chromosome 11 (markers between 12 Mb to 30 Mb) in this study was overlapped with this study. Moreover, in another F_2_ population (ICPL85063 × ICPL87119), a QTL was identified having marker interval from 12.6 Mb to 21.9 Mb with 56.45% PVE and 22.11 LOD score. This whole region exactly overlapped with the region identified in our study on chromosome 11 (markers between 12 Mb to 30 Mb). One minor QTL was identified from this study on chromosome 01, having marker interval between 2.8 Mb to 4.2 Mb, with 6.55% PVE and LOD 2.5. The region identified in our study on chromosome 1 (3171758bp) overlapped with this QTL region [64].

Two QTLs were identified at 4.0 cM and 14.6 cM on chromosome 9 for the Patancheru variant of SMD in F2 population ICP8863 × ICPL20096 [65]. We have found the physical position for qSMD1 nearby 28 Mb and for qSMD2 nearby 26 Mb. In our study, we have identified a strong, significant resistant region nearby 24 Mb on chromosome 9 for Bangalore race of SMD virus. The Bangalore race is highly severe and responsible for maximum pigeonpea yield losses for nearly all areas [66].

## 5. Conclusion

This study identified candidate genes, markers (SNPs and InDels), and haplotypes for FW and SMD resistance in pigeonpea using the GWAS and post-GWAS-analysis approaches. Key disease-resistance-related candidate genes were identified for both the disease and were significantly represented in the disease resistance pathway. Moreover, we have also compared the genomic regions and resistance donors identified in this study with the previous studies. Allelic distribution of identified markers was shown in resistant, moderately resistant, and susceptible genotypes. We have validated 12 KASP markers for FW and 4 KASP markers for SMD. We have also identified InDels from the significant regions. The haplotypes were identified from the significant regions for FW and SMD. H4 (TGT) from the *Cc10_4438493* region and H3 (CTTTTT) from *Cc11_21637656* are identified as a resistant haplotype for FW. For SMD, a resistant haplotype (CCCT) from Cc09_24131177 was identified.

In summary, this study has provided the promising regions and genes for FW on chromosomes 05, 10, and 11, and for SMD on chromosome 9. Two key resistant genes, viz., *Two component response regulator gene* (*Cc_11098*) and *Dof-type zinc finger DNA-binding family protein* (*Cc_23564*) were identified for FW resistance in pigeonpea. The most important resistant genes, viz., *LRR and NB-ARC domain disease resistance protein* (*Cc_21069*), *Flavonoid 3’, 5’-hydroxylase* (*Cc_21070*), and *Probable disease resistance protein* (*Cc_21071*) were identified for SMD resistance in pigeonpea. For FW, *Cc_10507586* region from chromosome 5 is identified in ICPL87119 and ICPL20096 donors. A region *Cc_21637656* from chromosome 11 is identified in the ICPL87119 but not in the ICPL20096. Similarly, a region *Cc_4438493* from chromosome 10 is identified in the ICPL20096 but not in the ICPL87119. For SMD, *Cc_24131177* region is identified in both the SMD resistance donors (ICPL20096 and ICPL87119). *Ccsmd04* region from chromosome 4 has been identified in ICPL87119 but not in the ICPL20096. These genomic regions can be introgressed and brought together into one line with the help of the markers identified in this study. The identified candidate genes will be helpful for future research on gene cloning and functional validation of causal mutations or genes responsible for the viral disease resistance in pigeonpea and other legume species.

## Data availability

All the data generated in the present study is provided in the Supplementary Information and sequencing data deposited as Bioproject PRJNA575817, PRJNA383013, PRJNA57990 in NCBI.

## Author Contribution

**Sagar Krushnaji Rangari:** Writing-original draft; investigation; methodology; formal analysis; validation**. Suma Krishnappa:** Data generation and data curation; writing-review and editing. **Mamta Sharma**: Writing-review and editing; data generation and data curation**. Vinay K Sharma**: Writing-review and editing, supervision. **Namita Dube**: Formal analysis; software; data curation; writing-review and editing. **Sunil S Gangurde**: Writing-review and editing; formal analysis. **Vinay Sharma**: Writing-review and editing. **Ragavendran Abbai**: Formal analysis; writing-review and editing. **Naresh Bomma:** Data generation and data curation. **Shruthi Belliappa:** Writing-review and editing. **K. L. Bhutia**: Writing-review and editing. **Mahendar Thudi**: Writing-review and editing. **Prakash I Gangashetty**: Writing-review and editing. **Manish K Pandey**: Conceptualization; data generation; supervision; methodology; funding acquisition, investigation, writing-original draft, writing-review and editing.

