## Supplementary figure 1 for "Mapping Fusarium Wilt and Sterility Mosaic Disease Resistance-Associated Genomic Regions and Haplotype Variants in Pigeonpea"

Supplementary figures

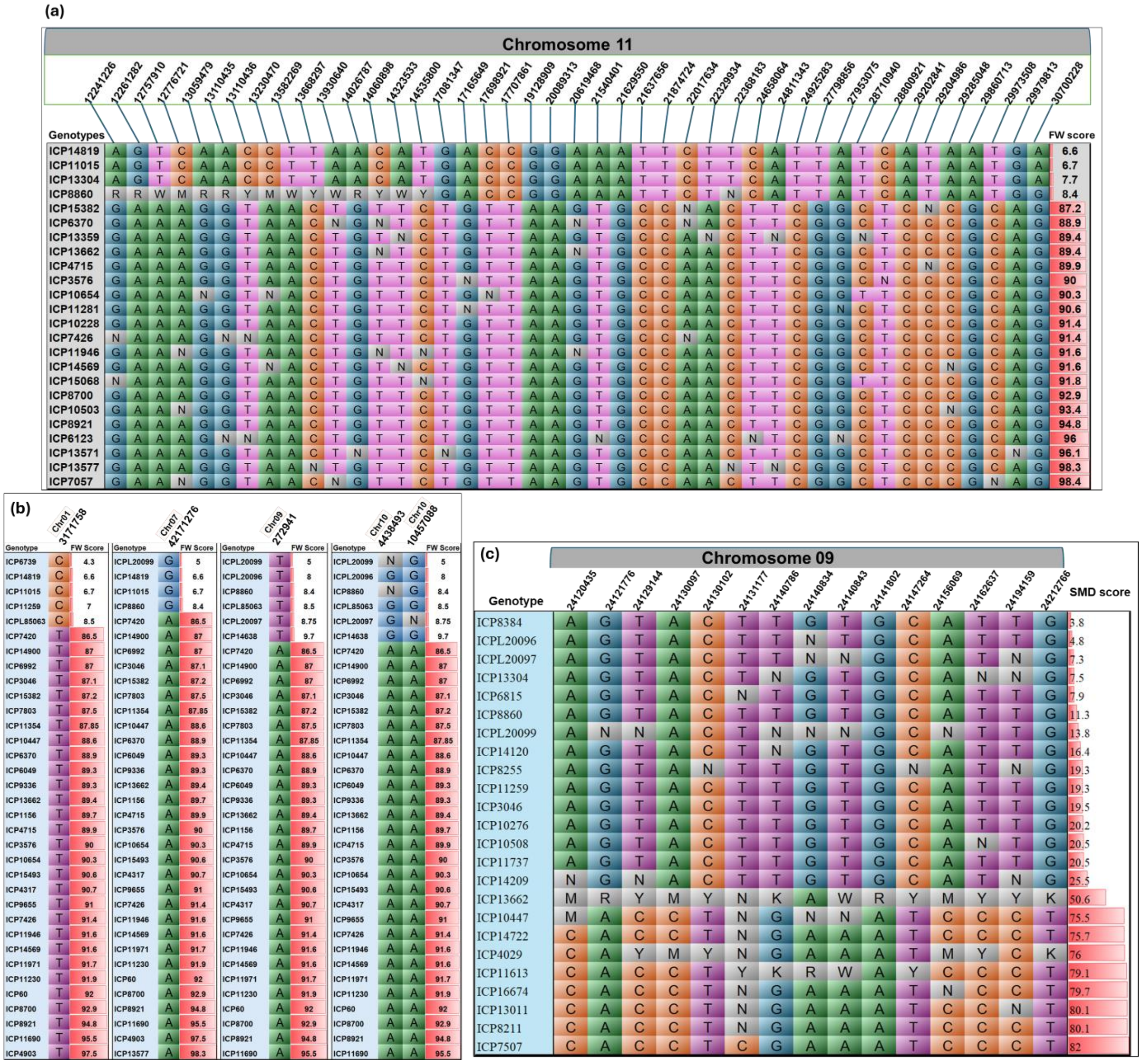


Supplementary Figure 1: Phenotype-wise allelic distribution of MTAs. The phenotypic scores of genotypes are represented at right side of each diagram in bar plot format (red gradient scale). In Grey colour missing and heterozygous alleles are represented (a) 43 MTAs on chromosome 11 showing allelic discrimination between resistant and susceptible genotypes. Four FW resistant and 20 susceptible genotypes are showing the favourable and unfavourable alleles respectively. The position of each marker is given in the green colour box. (b) One MTA on chromosome 01 have five resistant and 29 susceptible genotypes are showing the favourable and unfavourable alleles respectively. One MTA on chromosome 07 have four FW resistant and 30 susceptible genotypes are showing the favourable and unfavourable alleles respectively. One MTA on chromosome 09 have six FW resistant and 31 susceptible genotypes are showing the favourable and unfavourable alleles respectively. Two MTAs on chromosome 10 have six FW resistant and 31 susceptible genotypes are showing the favourable and unfavourable alleles respectively. (c) For SMD, six resistant, eight moderately resistant, and nine susceptible genotypes on chromosome 09 showing the favourable and unfavourable alleles respectively.
